# GPCR Antagonism via Antibody-Mediated Rewiring of Receptor Trafficking and Degradation

**DOI:** 10.1101/2025.08.08.669350

**Authors:** Kaitlin Rhee, Lawrence Shue, Akimasa Adachi, James Osei-Owusu, Dingjingyu Zhou, Aoxing Cheng, Apoorva Baluapuri, Edward P. Harvey, Karen Adelman, Meredith A Skiba, Jun R Huh, Andrew C Kruse, Xin Zhou

**Affiliations:** Department of Cancer Biology, Dana-Farber Cancer Institute, Boston, MA, USA; Department of Biological Chemistry and Molecular Pharmacology, Blavatnik Institute, Harvard Medical School, Boston, MA, USA; Department of Immunology, Blavatnik Institute, Harvard Medical School, Boston, MA, USA; The Eli and Edythe L. Broad Institute, Cambridge, MA, USA; Department of Biological Chemistry and Department of Pharmacology, University of Michigan School of Medicine, Ann Arbor, MI, USA

## Abstract

G-protein coupled receptors (GPCRs) represent one of the most important yet incompletely targeted classes of therapeutic proteins. Here, we report a novel strategy for functional GPCR antagonism through antibody-mediated endocytosis and lysosomal degradation. Our engineered bispecific antibodies, termed GPCR-TfR1 Targeting Chimeras (GTACs), achieve potent and selective downregulation of various GPCRs, including RXFP1 and CCR6, critical cancer and immune drug targets that are difficult to antagonize with conventional approaches. GTACs led to complete inhibition of receptor activity with over 100-fold greater potency than conventional antibody antagonists. Using four-color imaging, we elucidated the trafficking mechanism of both the target protein and TfR1 in living cells. The GTAC platform enables robust antagonism of signaling across diverse GPCR families and establishes induced endocytosis and lysosomal trafficking as a fundamentally new paradigm for therapeutic GPCR modulation.

## Introduction

G-protein coupled receptors (GPCRs) constitute the largest receptor class in humans with over 800 members, many of which play essential roles in human physiology and disease. Accordingly, roughly 1/3 of all FDA-approved drugs target GPCRs^1^. However, therapeutic development has only been successful for a subset of these receptors. Biogenic amine receptors comprise only 5% of all GPCRs, yet they account for 75% of approved GPCR-targeting drugs^2,3^. Peptide or protein-binding receptors comprise 15% of all GPCRs, yet they account for only 10% of GPCR-targeting drugs^3,4^. The vast majority of other receptors, including orphan receptors, lipid receptors, and those involved in sensory signaling, remain largely undrugged as well.

Current strategies to modulate GPCR signaling rely on competitive inhibition or stabilization of inactive conformations^5^ and are constrained by the challenge of developing selective high-affinity antagonists^6^, the presence of ligand-independent, constitutive signaling^7^, and high sequence homology within GPCR subfamilies that hinders selectivity^8^. These limitations are especially pronounced for GPCRs with protein or peptide ligands, which often exhibit tight ligand binding, autocrine or paracrine signaling, and high local ligand concentrations in the extracellular matrix^9^. Moreover, the discovery of functional biologic modulators against GPCRs has remained particularly difficult due to the limited accessibility of extracellular epitopes and the conformationally dynamic nature of these domains^6^. As a result, only a small number of antagonizing biologics targeting GPCRs have been developed despite the large number of disease-relevant targets in this family^1,10,11^.

In normal physiology, GPCR signaling is dynamically regulated via trafficking within the endolysosomal system. Internalization, recycling, and lysosomal degradation of prototypical GPCRs are influenced by extracellular ligand binding, intracellular phosphorylation by GPCR kinases, and β-arrestin recruitment^12–14^. Inspired by this natural mode of regulation, we hypothesized that induced proximity between a GPCR and an internalizing receptor would synthetically trigger receptor endocytosis and lysosomal degradation, effectively silencing GPCR signaling in a programmable and selective manner (**Fig. 1a**). We proposed this novel strategy for achieving pharmacological inhibition of GPCR activity at the cell surface.

**Figure 1.**
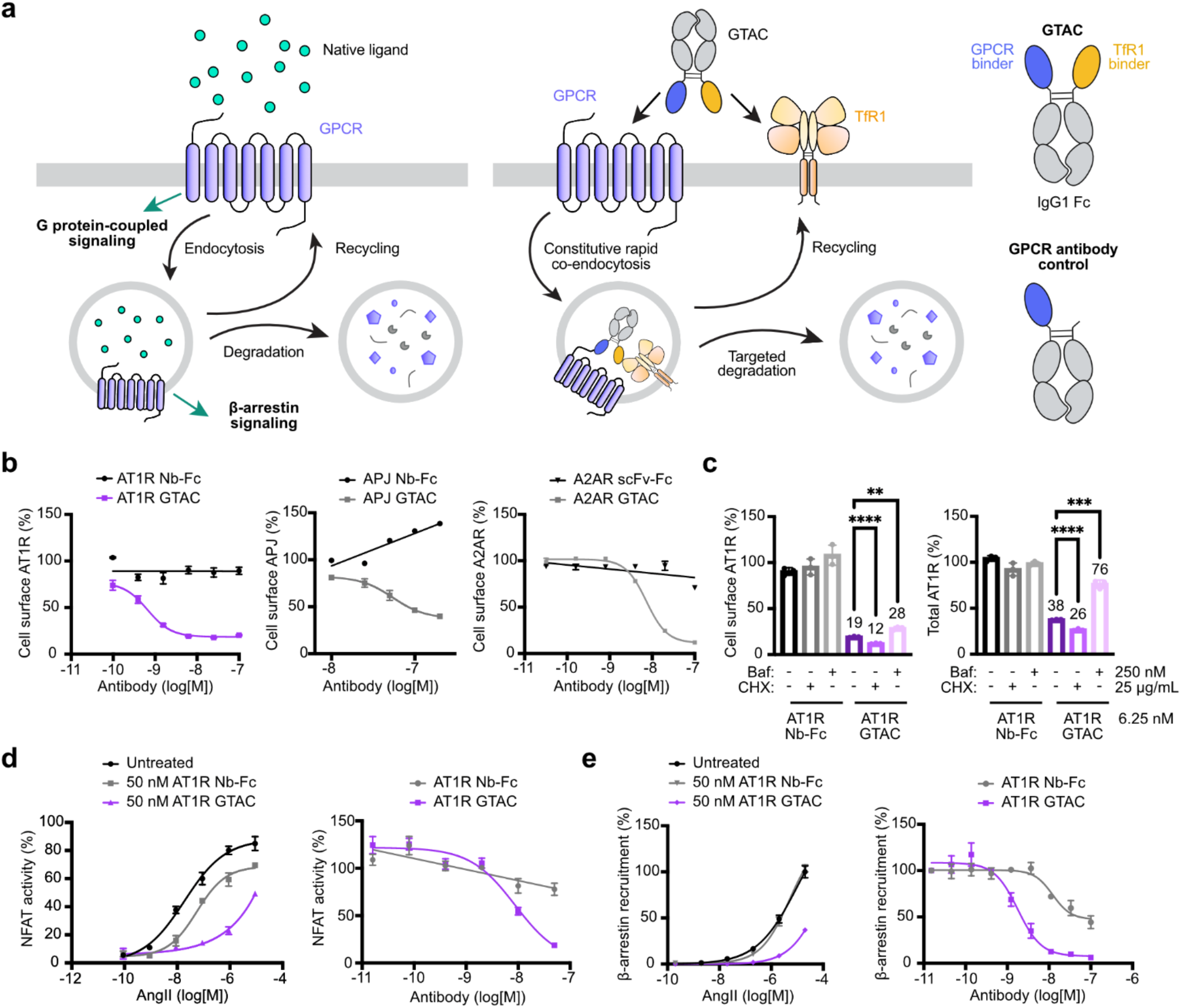
GTACs induce potent and sustained depletion of various GPCRs. **(a)** Schematic comparison of native versus engineered GPCR trafficking (left). Native agonists can induce GPCR internalization, followed by recycling or degradation via the endolysosomal pathway. GTACs direct surface GPCRs to degradation by leveraging rapid and constitutive TfR1-mediated internalization. Schematic of GTAC design (right), showing a bispecific antibody format comprising a GPCR-binding domain and a TfR1-binding domain fused to an IgG1 Fc region. The control antibody lacks the TfR1-binding domain. **(b)** GTAC-mediated depletion of cell surface GPCRs measured by anti-FLAG flow cytometry. Cell surface AT1R levels in FLAG-AT1R^+^ TfR1^+^ Expi293 cells after 6 h treatment with AT1R GTAC or Nb-Fc; cell surface APJ levels in FLAG-APJ^+^ TfR1^+^ HEK293T cells after 1 h treatment with APJ GTAC or Nb-Fc; and cell surface A2AR levels in FLAG-A2AR^+^ TfR1^+^ Expi293 cells after 6 h treatment with A2AR GTAC or scFv-Fc. Data represent mean ± standard deviation (s.d.) from n = 3 replicates for AT1R, n = 2 for APJ, and n = 3 for A2AR. **(c)** Cell surface AT1R levels (measured by anti-FLAG staining) and total AT1R levels (measured by GFP) in FLAG-AT1R-GFP^+^ TfR1^+^ Expi293 cells following 1.5 h pre-treatment with bafilomycin A1 (Baf) or cycloheximide (CHX), then 6 h treatment with AT1R GTAC or Nb-Fc. Data represent mean ± s.d. from n = 3 replicates. Statistical comparisons via unpaired two-tailed t tests for cell surface AT1R: No inhibitor vs. CHX: P < 0.0001; No inhibitor vs. Baf: P = 0.0013; for total At1R: No inhibitor vs. CHX: P < 0.0001; No inhibitor vs. Baf: P = 0.0001. **(d)** NFAT activation in AT1R^+^ TfR1^+^ NFAT-Luc^+^ Expi293 cells measured by luciferase activity. Cells were pre-treated for 2 h with 50 nM AT1R GTAC or Nb-Fc, followed by 4 h stimulation with varying concentrations of AngII (left), or pre-treated for 2 h with varying concentrations of AT1R GTAC or Nb-Fc and then stimulated for 4 h with 9 nM AngII. Data represent mean ± s.d. from n = 2 and n = 4 replicates, respectively. **(e)** β-arrestin recruitment in AT1R-Tango^+^ TfR1^+^ HTLA cells measured by luciferase activity after pre-treatment for 2 h with 50 nM AT1R GTAC or Nb-Fc, followed by 14 h activation with varying concentrations of AngII (left), or pre-treatment for 2 h with varying concentrations of AT1R GTAC or Nb-Fc followed by 14 h activation with 200 nM AngII (right). Data represent mean ± s.d. from n = 3 replicates for both experiments.

Here, we present the first example of extracellularly induced GPCR inhibition and degradation via bispecific antibody chimeras that couple GPCRs to transferrin receptor 1 (TfR1), a constitutively internalizing and recycling receptor. These antibodies, termed GPCR–TfR1 Targeting Chimeras (GTACs), enable potent functional antagonism of the relaxin family peptide receptor RXFP1, implicated in cancer progression, and the chemokine receptor CCR6, a driver of autoimmune T cell trafficking. RXFP1-targeting GTACs downregulate metastasis- and matrix remodeling–associated genes in an ovarian cancer cell model, while CCR6-targeting GTACs achieve complete blockade of chemotaxis in immune cells. Compared to conventional antagonists, GTACs exhibit more than two orders of magnitude greater potency and superior efficacy, underscoring the advantage of degradation-based antagonism.

We also demonstrate the generalizability of this strategy across three additional GPCRs from diverse families, including AT1R, APJ, and A2AR, and introduce a universal FLAG-targeting GTAC system to facilitate studies of GPCR trafficking and membrane- or endolysosomal localization– dependent signaling. Additionally, our work elucidates the mechanisms of GTAC function using a four-color live-cell imaging system and identifies protein engineering principles for designing degraders with high efficacy and specificity. These findings may have broad implications for antibody-based degrader development. Collectively, these studies establish proximity-induced endocytosis and lysosomal degradation as a powerful and modular paradigm for GPCR antagonism, with broad potential for applications in cancer, immune disorders, and a wide spectrum of GPCR-driven diseases.

## Results

### GTACs induce potent, TfR1-directed GPCR depletion, degradation, and signaling inhibition

GTACs are bispecific antibodies composed of a GPCR-binding domain and a TfR1-binding domain fused to an IgG1 Fc backbone, with variations in geometry and binder stoichiometry to achieve target-specific optimal activity (**Fig. 1a, SI Figs. 1–4**). TfR1 was chosen as the internalizing effector due to its broad tissue expression, rapid and constitutive internalization, and prior success in facilitating intracellular trafficking of other membrane protein families^15,16^. These features suggested the potential of TfR1 to serve as an effector for GPCR modulation across diverse cellular contexts. Moreover, its overexpression in tumor cells and activated immune cells makes it particularly relevant for cancer and immune modulation applications^16^.

We first constructed GTACs against three GPCRs with distinct ligand types and signaling and trafficking properties: the angiotensin II type 1 receptor (AT1R), which regulates blood pressure and cardiovascular function and has natural disease-associated mutations that dysregulate internalization^17,18,19^; the adenosine A2A receptor (A2AR), a critical immune regulator, where high extracellular adenosine concentrations (up to 100 μM) can limit conventional antagonist effectiveness, making degraders an appealing therapeutic strategy^20,21^; and the apelin receptor (APJ), which regulates cardiovascular function and binds multiple endogenous peptide isoforms, exhibiting ligand-biased signaling and complex trafficking behaviors^22–24^.

Receptors were tagged with an N-terminal FLAG epitope and overexpressed with TfR1 in HEK293T cells, or in some cases, in Expi293 cells to enhance expression. To target these GPCRs, we used antagonistic nanobodies (Nbs) or scFvs: Nb AT118-L for AT1R^25^, Nb JN241 for APJ^26^, and scFv Hu3F6 H5-L5 for A2AR (**Extended Data Fig. 1a, SI Figs. 1a, 1b, 2a**)^27^. Cell surface receptor levels were measured by anti-FLAG flow cytometry after treatment with GTACs or corresponding control antibodies lacking the TfR1 binder. GTACs efficiently reduced cell surface expression of all three GPCRs, achieving up to 75–80% reduction of AT1R within 6 h (IC_50_ = 1.3 nM), 90% reduction of A2AR within 6 h (IC_50_ = 7.3 nM), and 60% reduction of APJ within 2 h (IC_50_ = 39 nM) (**Fig. 1b**).

Receptor degradation was measured using FLAG-AT1R-GFP^+^ TfR1^+^ Expi293 cells. Treatment with AT1R GTAC induced up to 65% GFP downregulation (IC_50_ = 1.3 nM) (**Fig. 1c, Extended Data Fig. 1b, 1c**). Time-course analyses showed that GTAC activity was rapid and sustained, with maximal cell surface depletion reached within 1 h and maximal degradation by 6 h. These effects remained stable over 24 h, the longest time point measured (**Extended Data Fig. 1d**). Inhibiting lysosomal acidification with bafilomycin A1 significantly increased total AT1R levels from 38% to 76%, indicating degradation was mediated by lysosomes (**Fig. 1c, Extended Data Fig. 1c**). Bafilomycin had a smaller effect on cell surface AT1R levels, which rose modestly from 19% to 28%, suggesting that GTACs can still efficiently remove receptors from the surface even when lysosomal function is blocked (**Fig. 1c, Extended Data Fig. 1c**). Blocking protein synthesis with cycloheximide further enhanced GTAC efficacy, reducing total AT1R levels from 38% to 26% and cell surface levels from 19% to 12%. These results indicate that the overall extent of receptor loss reflects the combined effects of lysosomal GPCR degradation and new receptor synthesis.

Beyond depleting and degrading receptors, GTACs potently antagonize GPCR signaling. In functional assays, AT1R GTACs strongly suppressed angiotensin II (AngII)-induced NFAT reporter activation and β-arrestin recruitment (**Fig. 1d, 1e**). Compared to the antagonist Nb-Fc control, GTACs showed 10–100-fold higher potency and significantly greater maximal inhibition.

### GPCR modulation by FLAG-targeting GTAC

FLAG-tagged receptors are widely used in GPCR research. We leveraged the anti-FLAG M2 fragment antigen-binding domain (Fab) to develop FLAG-targeting GTACs, providing an orthogonal targeting strategy that does not interfere with endogenous ligand binding (**Fig. 2a, Extended Data Fig. 1a**). In FLAG-HA-AT1R-GFP^+^ TfR1^+^ Expi293 cells, FLAG GTAC induced potent depletion of both cell surface and total AT1R similar to the AT1R GTAC, achieving up to 75% internalization as measured by anti-HA flow cytometry and a 60% reduction in total receptor levels measured by GFP (IC_50_ = 600 pM) (**Fig. 2b**). Receptor depletion was both rapid and sustained, with over 50% of the maximal effect observed within 1 h and nearly 100% by 6 h, persisting over 24 h. FLAG-GTAC also significantly inhibited AngII-induced β-arrestin recruitment (**Fig. 2b**) and led to persistent cell surface depletion and receptor degradation (**Fig. 2c**). By contrast, 1 μM native ligand AngII triggered only transient AT1R internalization (∼50%) without leading to changes in total receptor levels (**Fig. 2c, 2d, Extended Data Fig. 1e**).

**Figure 2.**
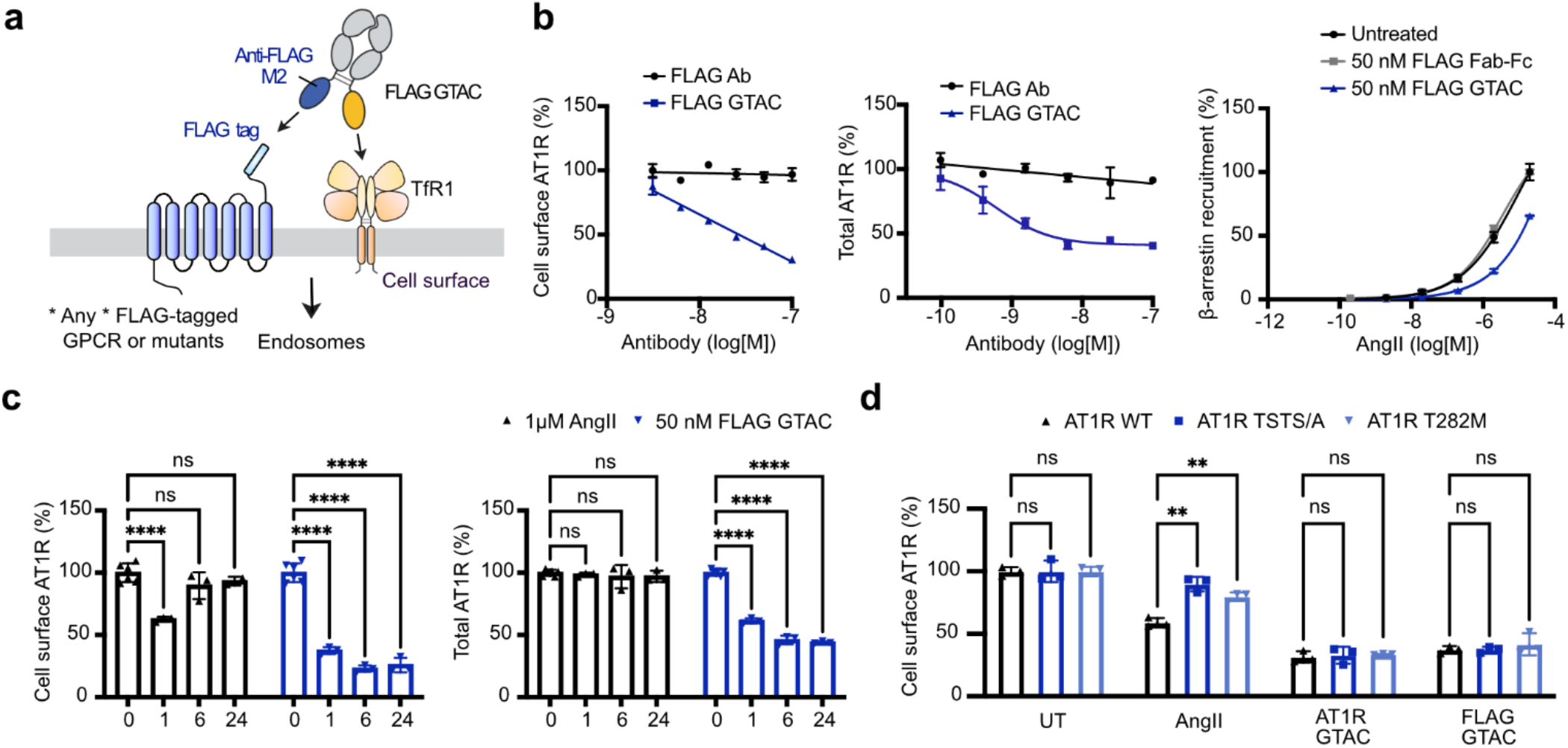
FLAG GTAC as a generalizable tool for GPCR modulation. **(a)** Schematic of FLAG-targeting GTACs for internalization and degradation of FLAG-tagged GPCRs. **(b)** Cell surface AT1R levels (left, measured by anti-HA staining) in FLAG-HA-AT1R^+^ TfR1^+^ Expi293 cells after 2 h treatment with FLAG GTAC or FLAG Fab-Fc; total AT1R levels (middle, measured by GFP fluorescence) in FLAG-AT1R-GFP^+^ TfR1^+^ Expi293 cells after 6 h treatment, measured by flow cytometry; and β-arrestin recruitment (right) in AT1R-Tango^+^ TfR1^+^ HTLA cells measured by luciferase activity after 2 h pre-treatment with 50 nM FLAG GTAC or Fab-Fc, followed by 14 h stimulation with AngII. Data represent mean ± s.d. from n = 3 replicates. **(c)** Cell surface AT1R levels (left, measured by anti-HA) and total AT1R levels (right, measured by GFP) over 24 h in FLAG-HA-AT1R-GFP^+^ TfR1^+^ Expi293 cells after treatment with either 1 μM AngII or 50 nM FLAG GTAC. Statistical comparisons via unpaired two-tailed t tests for cell surface AT1R experiments: AngII 0 h vs. 1 h, P < 0.0001; AngII 0 h vs. 6 h, P = 0.13; AngII 0 h vs. 24 h, P = 0.28; FLAG GTAC 0 h vs. 1 h, 6 h, or 24 h, P < 0.0001; for total AT1R experiments: AngII 0 h vs. 1 h, P = 0.38; AngII 0 h vs. 6 h, P = 0.45; AngII 0 h vs. 24 h, P = 0.24; FLAG GTAC 0 h vs. 1 h, 6 h, or 24 h, P < 0.0001. Data represent mean ± s.d. from n = 2-6 replicates. **(d)** Cell surface AT1R levels (measured by anti-HA) in TfR1^+^ Expi293 cells expressing FLAG-HA-AT1R-GFP with AT1R wild-type (WT), T282M mutant, or TSTS/A mutant. Cells were treated for 1 h with 1 μM AngII, 10 nM AT1R GTAC, or 50 nM FLAG GTAC. Statistical comparisons via unpaired two-tailed t tests for untreated conditions: WT vs. T282M or TSTS/A, P > 0.99; for AngII treatments: WT vs. T282M, P = 0.0021; WT vs. TSTS/A, P = 0.0015; for AT1R GTAC treatments: WT vs. T282M, P = 0.48; WT vs. TSTS/A, P = 0.77; for FLAG GTAC treatments: WT vs. T282M, P = 0.48; WT vs. TSTS/A, P = 0.77. Data represent mean ± s.d. from n = 3 replicates.

We further tested FLAG-GTACs on AT1R receptors harboring the T282M mutation^19^ or the T332A/S335A/T336A/S338A (TSTS/A) quadruple mutation^28,29^. These mutations are known to impair β-arrestin recruitment and ligand-induced internalization, either allosterically (T282M) or by eliminating key phosphorylation sites in the C-terminal tail (TSTS/A). Consistent with previous findings, AngII-induced internalization was significantly reduced in these variants (**Fig. 2d**). In contrast, both FLAG GTAC and AT1R GTAC efficiently depleted and degraded all AT1R mutants, with no significant differences observed in GTAC-mediated downregulation of WT versus mutant AT1R (**Fig. 2d**). These findings indicate that FLAG GTACs can reduce receptor levels independently of endogenous signaling pathways and suggest a possible approach for investigating post-internalization trafficking and signaling effects of GPCR mutations.

Overall, our studies with AT1R, APJ, and A2AR demonstrate that GTACs can mediate TfR1-dependent depletion, degradation, and signaling inhibition of various GPCRs and their mutants. Additionally, they can be modularly designed with binders in diverse formats, including Nbs, scFvs, and Fabs, or directed against functionally inert epitopes such as FLAG.

### RXFP1 GTACs achieve picomolar antagonistic potency and functional inhibition in ovarian cancer cells

Having established the broad application of GTACs across multiple GPCR families, we next focused on the therapeutic targeting of two challenging-to-drug receptors, RXFP1 and CCR6. RXFP1 is a GPCR activated by the peptide hormone relaxin-2 (RLN2), which normally mediates physiological adaptations of the renal and cardiovascular systems during pregnancy^30^. However, aberrant signaling through the relaxin family peptide receptors has been implicated in cancer growth, metastasis, and drug resistance^31–33^. Ovarian cancer, the most lethal gynecologic malignancy and a leading cause of cancer deaths in women, is often diagnosed at advanced stages, with resistance to standard chemotherapy and PARP inhibitors contributing to a five-year survival rate of less than 30%^34,35^. A recent shRNA screen of over 370 GPCRs identified RXFP1 as a key driver of ovarian cancer growth, metastasis, and cisplatin resistance through regulation of the Wnt, Notch, and ECM remodeling pathways, suggesting that RXFP1 antagonism could be a promising therapeutic strategy to inhibit ovarian cancer progression and metastasis^33,36–38^.

Despite this promise, RXFP1 remains exceptionally difficult to inhibit. This receptor exhibits picomolar affinity for RLN2, functions through autocrine signaling in cancer cells, and assembles a constitutive signaling complex that signals through a mechanism of autoinhibition and is highly sensitive to low ligand concentrations^33,39–41^. Previously described RLN2 analog inhibitors exhibit only modest affinity and limited efficacy, and no RXFP1 antagonists have advanced to clinical development^42,43^.

To develop an RXFP1 inhibitor, we first generated a Nb against RXFP1. Yeast surface display was used to screen and obtain a clone that binds competitively with RLN2 (**Fig. 3a**). The initial Nb exhibited an on-cell binding EC_50_ of 50 nM (**Extended Data Fig. 2a**). However, it showed limited functional antagonism in RLN2-induced cAMP signaling assays, indicating that it was insufficient to block receptor activation effectively, perhaps in part due to the high binding affinity and sensitivity to RLN2. To overcome this limitation, we performed affinity maturation via error-prone PCR and additional yeast selections, yielding a higher-affinity Nb with an on-cell binding EC_50_ of 1.8 nM (**Extended Data Fig. 2a**). This Nb exhibited enhanced competition with RLN2 in a flow-cytometry displacement assay, with a calculated Ki of 340 pM (**Fig. 3b**). Despite the improved binding, signaling inhibition by the Nb or Nb-Fc fusions remained modest, with significant inhibition observed only at micromolar concentrations of nanobody (**Extended Data Fig. 2b, 2d, 2f**).

**Figure 3.**
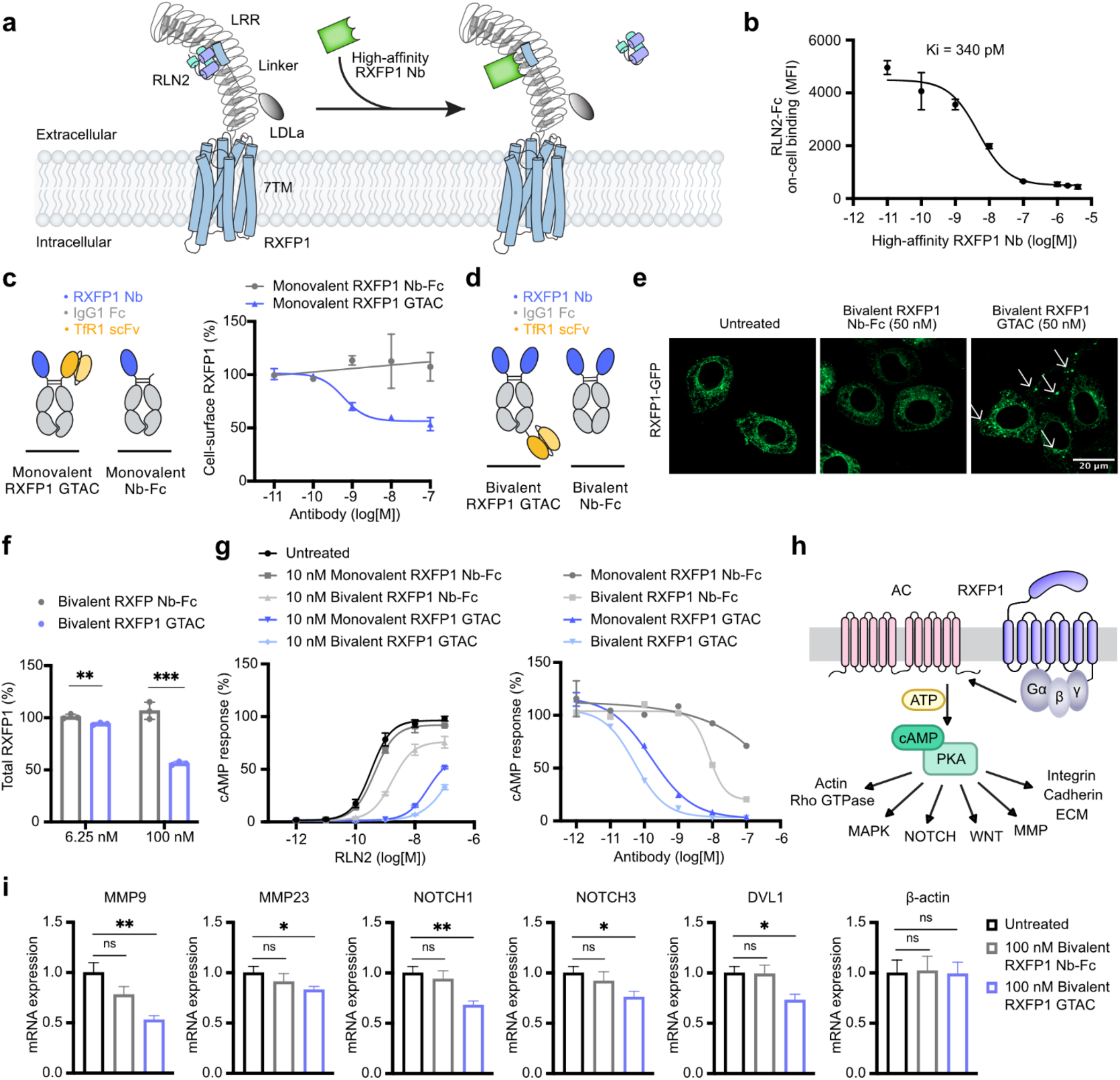
RXFP1-targeting GTACs exhibit ∼100-fold greater potency than a picomolar-affinity Nb antagonist and suppress downstream transcriptional programs in ovarian cancer cells. **(a)** Schematic of competition between high-affinity RXFP1 Nb and the native ligand RLN2 for RXFP1 binding. **(b)** Flow cytometry competition assay showing RLN2-Fc binding to Expi293 cells transfected with RXFP1 or empty vector, titrated with high-affinity RXFP1 Nb. Data represent mean ± s.d. from n = 3 replicates. **(c)** Schematic of monovalent RXFP1 GTAC and control antibody constructs (left). Cell surface RXFP1 levels (right) in FLAG-RXFP1^+^ TfR1^+^ Expi293 cells after 2 h treatment with RXFP1 GTAC or Nb-Fc. Data represent mean ± s.d. from n = 2 replicates. **(d)** Schematic of the optimized bivalent RXFP1 GTAC with highest potency and the corresponding control antibody. **(e)** Confocal microscopy images showing RXFP1-GFP localization in FLAG-RXFP1-GFP^+^ TfR1^+^ Hela cells after 6 h treatment with 50 nM bivalent RXFP1 GTAC, Nb-Fc, or untreated controls. Arrows indicate intracellular puncta. Scale bar: 20 μm. **(f)** Total RXFP1 levels (measured by GFP fluorescence) in FLAG-RXFP1-GFP^+^ TfR1^+^ Expi293 cells after 6 h treatment with bivalent RXFP1 GTAC or Nb-Fc. Statistical comparisons via unpaired two-tailed t tests at 6.25 nM: P = 0.009; at 100 nM: P = 0.0005. Data represent mean ± s.d. from n = 3 replicates. **(g)** RLN2-induced cAMP signaling measured in RXFP1^+^ TfR1^+^ GloSensor^+^ HEK293T cells using a split-luciferase cAMP assay. Cells were pre-treated for 2 h with 10 nM RXFP1 GTAC or Nb-Fc, followed by 30 min stimulation with RLN2 at varying concentrations (left), or pre-treated with RXFP1 GTAC or Nb-Fc at varying concentrations, followed by stimulation with 1 nM RLN2 (right). Data represent mean ± s.d. from n = 2 replicates (left) or n = 1 replicate (right). **(h)** Schematic of downstream RXFP1–cAMP signaling pathways implicated in tumor progression, including activation of PKA-dependent transcriptional programs involved in ECM remodeling, cell adhesion, and metastasis. **(i)** Quantitative PCR measurements of RXFP1-regulated genes in OVCAR-8 ovarian cancer cells. Cells were serum-starved, untreated or treated overnight with 100 nM bivalent GTAC or Nb-Fc, then stimulated with 10 nM RLN2 for 8 h. mRNA levels of *MMP9, MMP23B, NOTCH1, NOTCH3, DVL1*, and *β-actin* were measured by qPCR and normalized to GAPDH, with the value for untreated cells set to 1. *β-actin* is shown as a control. Statistical comparisons of Untreated vs. GTAC conditions via unpaired two-tailed t tests for *MMP9*: P = 0.0057; for *MMP23:* P = 0.039; for *NOTCH1*: P = 0.0066; for *NOTCH3*: P = 0.024, for *DVL1*: P = 0.017; for *β-actin*: P = 0.96. Data represent mean ± s.d. from n = 3 replicates.

Using a doxycycline-inducible FLAG-RXFP1^+^ HEK293T cell line, we validated that cell surface expression levels positively correlate with RLN2-induced RXFP1 cAMP signaling (R^2^ = 0.97, **Extended Data Fig. 2c**), supporting the rationale for the GTAC approach to antagonize RXFP1. We generated an RXFP1 GTAC using a knob-into-hole Fc containing one high-affinity RXFP1 Nb and one anti-TfR1 H7 scFv. RXFP1 GTAC led to up to 44% reduction of surface RXFP1 with an IC_50_ of 594 pM, whereas the control monovalent Nb-Fc induced no cell surface RXFP1 reduction, indicating TfR1-mediated RXFP1 internalization (**Fig. 3c**).

We further constructed a panel of RXFP1 GTACs varying in geometry, binding stoichiometry, and incorporating either low- or high-affinity RXFP1 Nbs (**SI Fig. 3**). From this panel, we identified an optimized bivalent RXFP1 GTAC design, comprising two high-affinity RXFP1 Nbs fused to the Fc N-terminus and one TfR1 scFv at the C-terminus (**Fig. 3d**). In FLAG-RXFP1^+^ TfR1^+^ Expi293 cells, the bivalent RXFP1 GTAC achieved a maximum 45% depletion of cell surface receptors, with an IC_50_ of 190 pM. Fluorescence imaging in FLAG-RXFP1-GFP^+^ TfR1^+^ HeLa cells showed that untreated cells exhibited predominantly perinuclear localization of RXFP1-GFP, consistent with prior reports of ER and Golgi accumulation (**Fig. 3e**) ^44^. Upon GTAC treatment, RXFP1 redistributed to punctate intracellular vesicles, suggesting that GTACs redirect the membrane pool into the endolysosomal pathway. The large intracellular pool of RXFP1 and potential shuttling of RXFP1 from intracellular compartments to the membrane may explain why only partial depletion of RXFP1 was achieved with GTAC. In FLAG-RXFP1-GFP^+^ TfR1^+^ HEK293T cells, GTAC treatment achieved up to ∼50% reduction in total receptor levels (**Fig. 3f**).

Importantly, RXFP1 GTACs demonstrated dramatically enhanced functional antagonism. In cAMP GloSensor assays with FLAG-RXFP1^+^ TfR1^+^ HEK293T cells, both the monovalent and bivalent RXFP1 GTACs achieved nearly 100% maximal inhibition of RLN2-induced signaling, whereas the monovalent Nb-Fc control reached only 29% inhibition at 100 nM (**Fig. 3g, SI Table 2**). The bivalent Nb-Fc showed improved maximal signaling inhibition compared to the monovalent Nb-Fc, likely because it also moderately reduced cell surface RXFP1 levels (**Extended Data Fig. 2f**). Consistent with this, the bivalent GTAC was three-fold more potent than the monovalent GTAC in both RXFP1 depletion and cAMP inhibition. Moreover, the bivalent RXFP1 GTAC exhibited over 150-fold greater potency than the bivalent Nb-Fc (GTAC IC_50_ = 52 pM; Nb-Fc IC_50_ = 7.9 nM) (**Fig. 3g, Extended Data Fig. 2f, SI Table 2**). Comparisons of GTACs incorporating high-versus low-affinity Nb arms revealed similar receptor depletion but differing levels of signaling inhibition, suggesting that high-affinity binding contributes to antagonism, possibly through sustained receptor engagement or enhanced ligand competition (**Extended Data Fig. 2d, 2f**). Time-course experiments demonstrated that GTAC-mediated signaling inhibition persisted for over 24 h (**Extended Data Fig. 2e, 2g)**. These data demonstrate that RXFP1 GTACs, designed with a high-affinity RLN2-competitive Nb, can achieve complete and durable signaling antagonism and may represent the first reported RXFP1 biologic inhibitor with picomolar potency.

Unlike AT1R and many other GPCRs, RXFP1 is not known to undergo significant ligand-induced internalization; consistent with previous reports, we did not observe RLN2 to induce substantial downregulation of RXFP1 (**Extended Data Fig. 3c**)^45^. Our results highlight the ability of RXFP1 GTACs to reprogram the trafficking and signaling of a receptor with inherently low native internalization capacity. Because GTACs function through induced internalization, they may be less effective against cell-permeable agonists, such as AZD5462^46^. Indeed, in FLAG-RXFP1^+^ TfR1^+^ GloSensor^+^ HEK293T cells, pre-treatment with 10 nM bivalent RXFP1 GTAC for 2 h effectively antagonized RLN2-induced signaling, whereas AZD5462-induced signaling was minimally affected (**Extended Data Fig. 3a, 3b**).

To evaluate the therapeutic relevance of RXFP1-targeting GTACs, we characterized their activity in OVCAR-8, a cisplatin-resistant high-grade serous ovarian cancer cell line with endogenous RXFP1 and TfR1 expression. Flow cytometry confirmed that the RXFP1 Nb-Fc, TfR1 scFv, and GTAC all bound to OVCAR-8 cells in a dose-dependent manner, verifying RXFP1 and TfR1 expression (**Extended Data Fig. 3d**). We then examined the effect of GTAC treatment on downstream gene expression. The RLN2–RXFP1–cAMP axis regulates multiple pro-tumorigenic pathways, including extracellular matrix remodeling, Notch signaling, and Wnt signaling (**Fig. 3h**). Following overnight treatment with the bivalent RXFP1 GTAC and 8 h RLN2 stimulation, qPCR showed significant downregulation of key RXFP1 target genes in OVCAR-8 cells, including *MMP9* and *MMP23B* (ECM remodeling); *NOTCH1* and *NOTCH3* (Notch); and *DVL1* (Wnt) (**Fig. 3i**). In contrast, the control bivalent Nb-Fc had no significant effects.

Altogether, these results establish RXFP1 GTACs as highly potent, durable, and mechanistically distinct antagonists that overcome the limitations of traditional competitive antagonists by combining cell surface receptor depletion with ligand blockade. Moreover, RXFP1 GTAC effectively suppressed downstream transcriptional programs associated with tumor progression and metastasis, suggesting their potential as a novel therapeutic strategy for ovarian cancer.

### Developing a CCR6 GTAC for immune modulation

We next explored whether GTACs could be applied to immune cell targets. Chemokine receptors are attractive therapeutic targets in autoimmune diseases and chronic inflammatory disorders but remain challenging to inhibit due to competition with high local concentrations of endogenous chemokines, broad ligand-binding pockets, and the development of antagonist tolerance^47–49^. C-C chemokine receptor type 6 (CCR6), for example, mediates immune cell recruitment to inflamed tissues^50–54^, is expressed on subsets of T cells, B cells, neutrophils, and dendritic cells^55,56^, and represents a promising target in autoimmune and chronic inflammatory diseases such as psoriasis, rheumatoid arthritis, and ulcerative colitis^50,54,57^. Despite its importance, there are currently no approved drugs targeting CCR6, despite extensive drug discovery efforts. Monoclonal antibodies have shown incomplete receptor blockade^57,58^, while small-molecule inhibitors have limited selectivity because they also inhibit CCR7 and CXCR2, and further clinical studies are needed to determine their therapeutic potential^59^. These challenges underscore the need for alternative strategies to achieve complete and selective CCR6 inhibition.

In RXFP1 GTACs and other previously developed tumor antigen-targeting TfR1 degrader constructs, we employed a transferrin-competitive TfR1 binder called H7 to leverage the elevated expression of TfR1 in tumor cells for enhanced cancer targeting^16,60^. However, targeting GPCRs in non-cancer cells may require non-competitive transferrin binding to minimize impacts on cellular iron uptake. We therefore developed a second-generation GTAC design that incorporates a non-competitive anti-TfR1 Nb. Furthermore, we introduced a point mutation into the Nb to lower the affinity to TfR1 and therefore potentially facilitate TfR1 recycling. We constructed and expressed CCR6-targeting GTACs using an anti-CCR6 antibody Fab fragment combined with the anti-TfR1 Nb with either high or low affinity. CCR6 was overexpressed with an N-terminal FLAG tag in Jurkat cells, a human T lymphocyte cell line with high endogenous TfR1 expression, as the model system.

We evaluated the activity of CCR6-targeting GTAC variants in CCR6^+^ Jurkat T cells and compared their effects to those of an antagonist antibody control (**Fig. 4a**). Both the high and low affinity GTACs induced rapid and potent depletion of cell surface CCR6, with IC_50_ values of 1.7 nM and 1.8 nM, respectively, achieving approximately 70% reduction within 2 h and up to 85% cell surface depletion by 24 h, whereas the CCR6 Fab showed no effect (**Fig. 4b**). The high affinity CCR6 GTAC led to partial co-depletion of cell surface TfR1 within 2 h, while the low affinity mutants did not exhibit this effect (**Fig. 4b**). Western blot analysis confirmed that both the high affinity and the low-affinity GTACs achieved near-complete degradation of CCR6 after an 18 h incubation (**Fig. 4c**). Consistent with flow cytometry data, TfR1 levels remained stable following treatment with the low affinity GTAC, whereas reduced TfR1 expression was observed with the high affinity GTAC.

**Figure 4.**
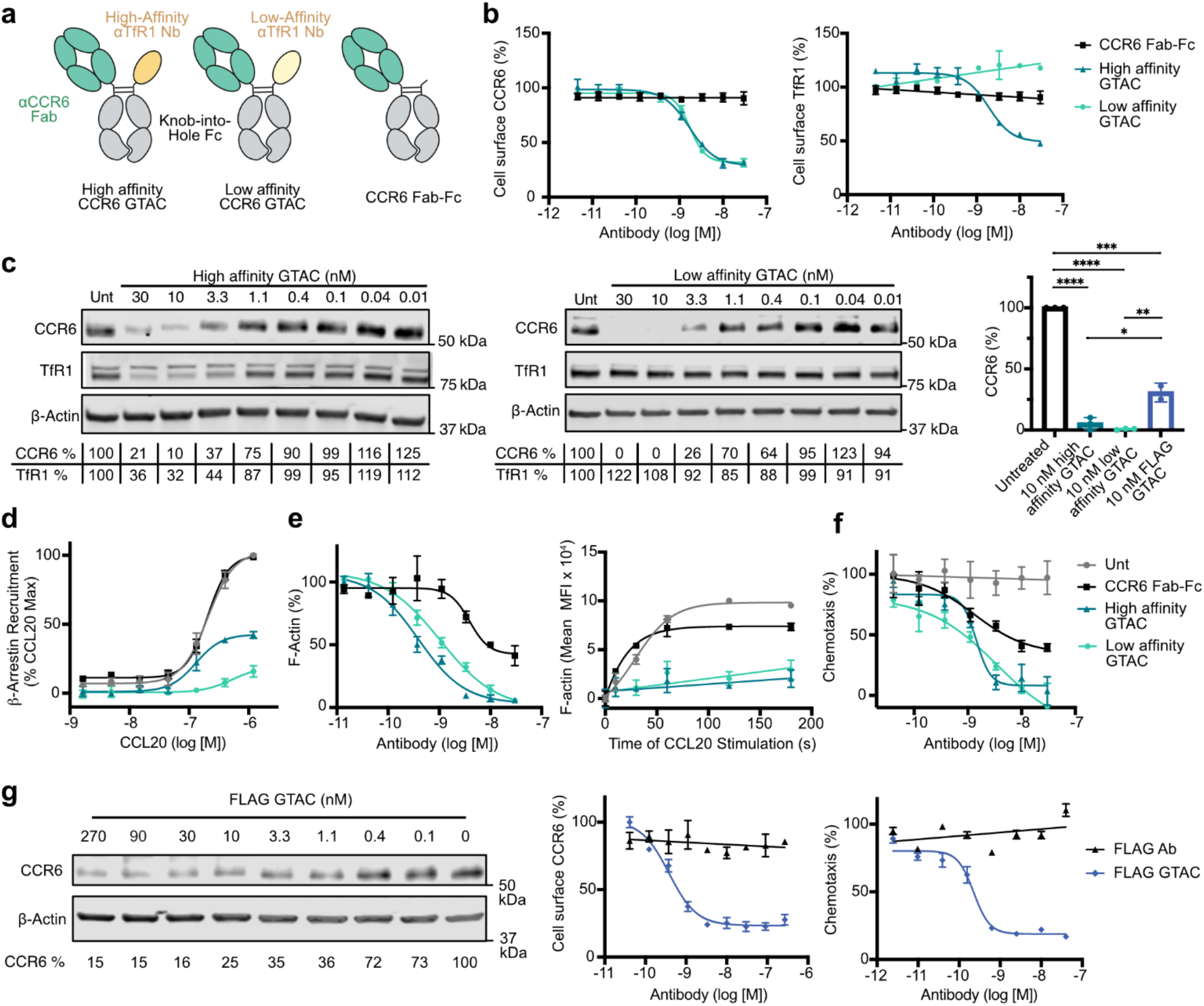
GTAC-mediated degradation and functional inhibition of CCR6. **(a)** Schematic of CCR6-targeting GTACs. Each GTAC comprises an anti-CCR6 Fab fused to an anti-TfR1 Nb via a knob-into-hole Fc backbone to promote heterodimerization. A CCR6 Fab-Fc lacking the TfR1-binding domain was included as a control. **(b)** Cell surface FLAG-CCR6 and TfR1 expression in CCR6^+^ Jurkat cells after 2 h treatment with GTACs that have either a high-affinity TfR1 binder, or low-affinity binder, measured by anti-FLAG or anti-TfR1 flow cytometry. Data represents mean ± s.d. from two replicates. **(c)** Western blot analysis and quantification of total CCR6 and TfR1 protein levels in CCR6^+^ Jurkat cells after 18 h GTAC treatment. Bar graphs of quantification of CCR6 levels following 10 nM GTAC treatment is shown at right. Statistical comparisons for: high affinity vs. Untreated: P < 0.0001; low affinity vs. Untreated: P < 0.0001; FLAG vs. Untreated: P = 0.0004; high affinity vs. FLAG: P = 0.02; low affinity vs. FLAG: P = 0.005 via unpaired two-tailed t-tests. Data are presented as mean from three replicates (two for FLAG). **(d)** β-arrestin recruitment measured via Tango assay in CCR6-Tango^+^ TfR1^+^ HTLA cells. Cells were pre-treated with 10 nM GTAC for 1 h, followed by overnight incubation with CCL20. Data are presented as mean ± s.d. from three replicates. **(e)** Flow cytometry quantification of total F-actin in CCR6^+^ Jurkat cells with fixation and phalloidin staining. Left: cells were treated with varying GTAC doses for 2 h, followed by 1 min stimulation with 50 nM CCL20. Right: cells were treated with 10 nM GTAC for 2 h and stimulated with 50 nM CCL20 for various durations. Data are presented as mean ± s.d. from two (left) or three (right) replicates. **(f)** Chemotaxis of CCR6^+^ luciferase^+^ Jurkat cells measured by Transwell migration assay after 2 h treatment & 2 h migration with CCL20 as the chemoattractant. Data are presented as mean ± s.d. from three replicates. **(g)** Western blot analysis of total CCR6 levels in FLAG-CCR6^+^ Jurkat cells after 18 h GTAC treatment (left). Flow cytometry analysis of cell surface FLAG-CCR6 expression in CCR6^+^ Jurkat cells after 2 h FLAG Ab/GTAC treatment (middle). Data represent mean ± standard deviation from two replicates. Chemotaxis of FLAG-CCR6^+^ luciferase^+^ Jurkat cells measured by Transwell migration assay after 2 h FLAG Ab/GTAC treatment & 2 h migration with CCL20 as the chemoattractant (right). Data represent mean ± standard deviation from three replicates.

Functionally, GTACs achieved complete suppression of CC chemokine ligand 20 (CCL20)-mediated CCR6 signaling and immune cell chemotaxis. In β-arrestin recruitment assays, 10 nM GTAC incubation for 18 h led to ∼100% signal inhibition up to 100 nM CCL20 and ∼80% inhibition at 1 μM CCL20, whereas 10 nM CCR6 Fab caused no inhibition (**Fig. 4d**). The low affinity GTAC demonstrated stronger inhibitory activity than high affinity GTAC, consistent with its enhanced receptor depletion. Activation of CCR6 initiates downstream F-actin-mediated cytoskeletal remodeling; in F-actin polymerization assays, 2 h GTAC treatment inhibited up to 100% of CCL20-induced actin polymerization after 1 min of ligand stimulation, with IC_50_ values of 0.39 nM for high affinity and 1.0 nM for low affinity GTACs, compared to only ∼55% maximal inhibition with the CCR6 Fab control (**Fig. 4e**). These effects were sustained across varying durations of CCL20 exposure.

Moreover, in Transwell chemotaxis assays, 2 h GTAC pretreatment at varying doses suppressed CCR6-mediated Jurkat cell migration by up to 100% with the low affinity GTAC and up to 90% with the high affinity GTAC, with IC_50_ values of 3.5 nM and 1.4 nM, respectively, whereas treatment with the CCR6 antibody alone resulted in up to ∼60% inhibition (**Fig. 4f**). The partial inhibition of CCL20-mediated chemotaxis by the antagonist antibody is consistent with previous reports^57,58^. These findings demonstrate that GTACs are highly potent in functionally inhibiting CCR6 and can overcome the limitations of incomplete inhibition observed with the antagonist antibody.

We next investigated whether the FLAG-targeting GTAC could result in similar antagonistic effects. Treatment of FLAG-CCR6^+^ Jurkat cells with a high-affinity TfR1 Nb-based FLAG GTAC resulted in up to ∼75% cell surface depletion after 2 h (IC_50_ = 380 pM), and potent chemotaxis inhibition of up to ∼85% (IC_50_ = 230 pM), showing comparable activities to the CCR6 GTAC, whereas the control FLAG antibody showed no depletion or inhibitory effect (**Fig. 4g**). These results align with findings for FLAG-GTAC modulation of AT1R, further demonstrating FLAG GTAC as a robust tool for regulating FLAG-tagged GPCR levels and signaling. Despite similar cell surface depletion levels and antagonism potency, the FLAG GTAC showed reduced degradation efficacy and potency (∼70% degradation at 10 nM) compared to the CCR6 GTACs (∼95% degradation at 10 nM), suggesting that more receptors remain trapped inside cells rather than being degraded (**Fig. 4c, 4g**). Together, these findings demonstrate that CCR6 GTACs achieve high-efficacy target depletion and degradation, effectively block signaling and function across multiple readouts, completely inhibit chemotaxis, and demonstrate superiority compared to a CCR6 antagonist antibody, suggesting the promise of GTACs as a novel therapeutic strategy to target chemokine receptors in autoimmune and inflammatory disorders.

### Four-color live-cell imaging reveal kinetics and trafficking features of GTACs

To elucidate the intracellular trafficking mechanisms of GTAC-mediated GPCR degradation, we developed a four-color live-cell imaging platform in HeLa cells. Cells were engineered to stably express CCR6-GFP and TfR1-BFP, as well as mCherry-tagged Rab5, Rab7, Rab11, or LAMP1, labeling early endosomes, late endosomes, recycling endosomes, and lysosomes, respectively (**Extended Data Fig. 4a–d**). GTACs were conjugated to Alexa Fluor 647. Cells were sorted via FACS for uniform marker expression. Using confocal microscopy, we tracked trafficking and degradation dynamics over time. Both the high and low affinity CCR6 GTACs induced near-complete loss of CCR6-GFP signals after 24 h, while only the high affinity GTAC also reduced TfR1-BFP levels, consistent with flow cytometry and western blot data (**Fig. 5a**). By contrast, treatment with the native ligand CCL20 induced internalization of CCR6-GFP but did not promote its degradation.

**Figure 5.**
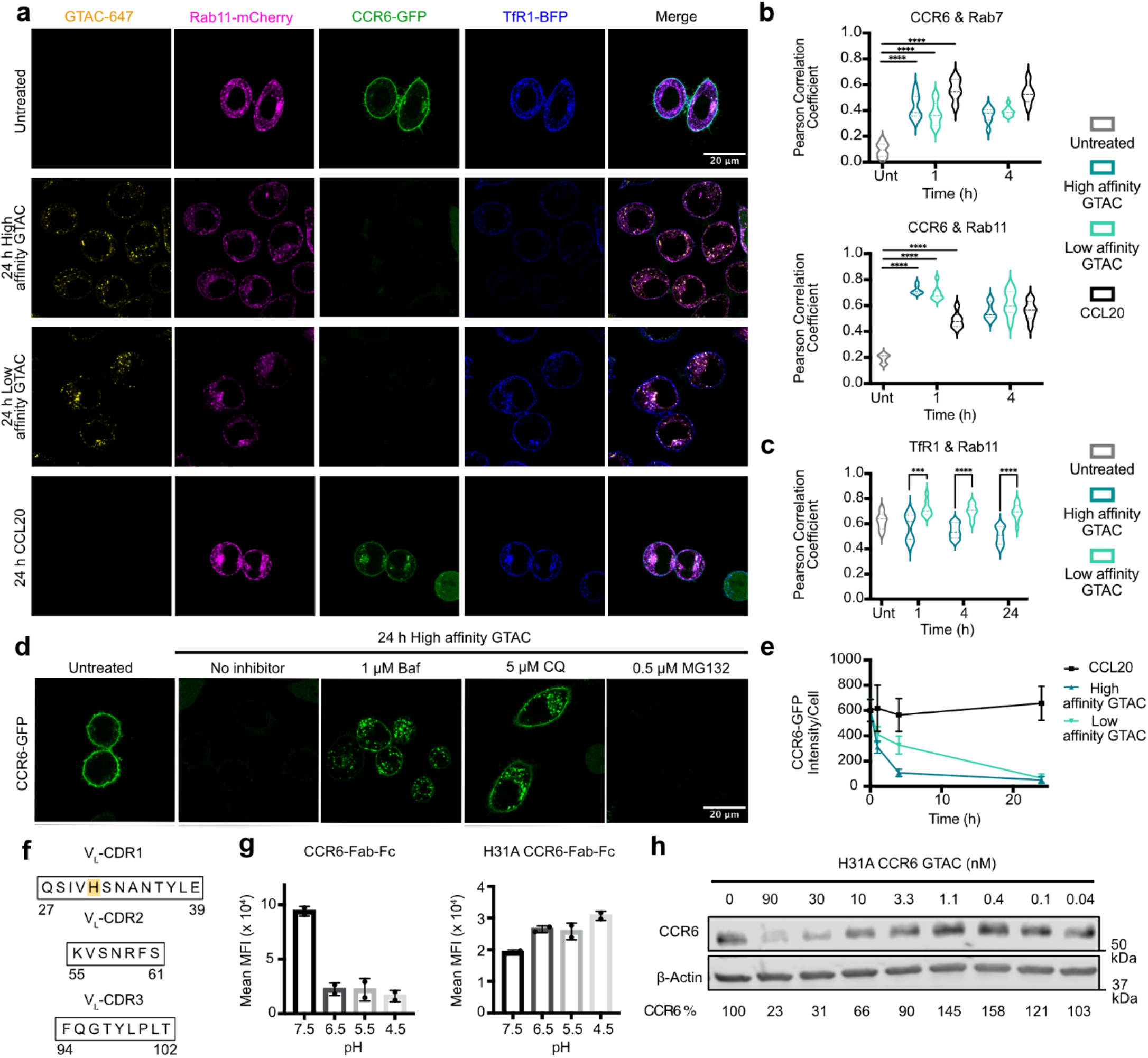
Elucidating the mechanisms of GTAC-mediated GPCR Degradation. **(a)** Multiplexed four-color live-cell confocal microscopy of HeLa cells expressing Rab11-mCherry, CCR6-GFP, and TfR1-BFP with 24 h treatment of 10 nM GTAC conjugated to Alexa Fluor 647 or 50 nM CCL20. **(b)** Pearson colocalization analysis of CCR6-GFP & Rab7-mCherry (late endosomes), CCR6-GFP and Rab11-mCherry (recycling endosomes), or **(c)** TfR1-BFP and Rab11-mCherry at different times with GTAC treatment. P < 0.0001 for all comparisons shown in (b). Low affinity GTAC induced higher TfR1-BFP colocalization with Rab11-mCherry at 1 (P = 0.0063), 4 (P = 0.000055), and 24 h (P = 0.000009). Data are presented as mean ± s.d. n of cells for analyses are summarized in SI Table 4. P values were determined by unpaired two-tailed t-tests. **(d)** Confocal microscopy of HeLa cells expressing CCR6-GFP with 24 h high affinity GTAC treatment in addition to the lysosomal inhibitors bafilomycin & chloroquine or the proteasome inhibitor MG132. **(e)** Quantification of CCR6-GFP in Rab5-mCherry^+^, CCR6-GFP^+^, TfR1-BFP^+^ HeLa cells by confocal microscopy with 1 h, 4 h, or 24 h treatment with GTAC variants or CCL20. Data are presented as mean ± s.d. n of cells for analyses are summarized in SI Table 4. **(f)** CDR regions of the αCCR6 Fab light chain. The Histidine 31 residue that is mutated in **(g)** and **(h)** is highlighted in yellow. **(g)** Flow cytometry showing cell surface binding of 5 nM CCR6-Fab-Fc or 125 nM H31A CCR6-Fab-Fc to CCR6^+^ and CCR6^−^ Jurkat cells after 10 min incubation on ice. Data represent mean ± s.d. from two replicates. **(h)** Western blot analysis and quantification of total CCR6 levels in CCR6^+^ Jurkat cells after 18 h H31A CCR6 GTAC treatment.

Colocalization analysis showed that following GTAC treatment, CCR6 exhibited increased colocalization with both Rab7 and Rab11, indicating enhanced trafficking through both late and recycling endosomal compartments (**Fig. 5b**). These results indicate that after GTAC-induced internalization, CCR6 does not exclusively traffic to lysosomes for degradation but instead enters both recycling and degradation pathways. Therefore, GTAC-mediated CCR6 degradation likely results from accelerated receptor internalization, while CCR6 continues to be sorted between recycling and degradation pathways. Whether the proportions of sorting to recycling and degradation change with GTAC treatment remains to be determined. Additionally, the low affinity GTAC resulted in increased colocalization of TfR1 with Rab11 compared to the high affinity GTAC, consistent with our hypothesis that reduced TfR1 affinity facilitates TfR1 recycling (**Fig. 5c**).

To determine the degradation pathway responsible for CCR6 loss, cells were pretreated for 1 h with lysosomal inhibitors (bafilomycin A1 or chloroquine) or the proteasome inhibitor MG132, followed by an 18 h incubation with GTACs. Lysosomal inhibition completely blocked GTAC-induced CCR6-GFP degradation, whereas proteasome inhibition had no effect, confirming that GTACs drive lysosome-dependent, not proteasome-dependent, degradation (**Fig. 5d**). CCR6 degradation was observed as early as 1 h, although the low affinity GTAC required more time to achieve maximal degradation; both variants ultimately led to near complete CCR6 degradation at 24 h (**Fig. 5e**).

Lastly, the observation that CCR6 underwent nearly complete degradation, a level higher than that seen with most other GPCRs tested, prompted further characterization of the CCR6 binder. Interestingly, sequence analysis of the CCR6 Fab revealed a histidine residue (H31) within the CDR1 region of its light chain (**Fig. 5f**), suggesting potential pH sensitivity. Indeed, flow cytometry measurements showed that CCR6 Fab binding to CCR6^+^ Jurkat cells was significantly reduced at acidic pH values (6.5, 5.5, and 4.5) compared to pH 7.5 (**Fig. 5g**).

To probe the role of this pH sensitivity, we mutated H31 to alanine (H31A), which abolished the pH-dependent binding, as evidenced by stable binding across lower pH values, though overall binding was reduced (**Fig. 5g**). We then tested the H31A CCR6 GTAC with the high affinity TfR1 Nb in FLAG-CCR6^+^ Jurkat cells. While it retained the ability to degrade CCR6, its potency and efficacy were reduced compared to the unmutated CCR6 GTAC, achieving only 34% degradation at 10 nM versus 90% with the original GTAC (**Fig. 5h, 4c**). The decrease in degradation may result from the combined effects of loss of pH-dependent dissociation and the overall reduction in CCR6 binding. Taken together with our earlier observations that pH-insensitive FLAG and AT1R GTACs, achieved similarly high receptor depletion and signaling inhibition while leading to lower degradation levels, these findings suggest that pH-dependent binding is not necessary for GTAC-induced antagonism. However, pH-dependent binding may enhance degradation, possibly by facilitating decoupling of the receptor from recycling TfR1 in early endosomes, and its role in influencing GTAC activities across different targets merits further investigation in future studies.

We further investigated how geometry, affinity, and avidity influence GTAC activity for AT1R, RXFP1, and CCR6 (**SI Fig. 2–4; SI Tables 1–3**). Overall, despite differences in potency and degradation efficacy, most GTACs achieved substantial receptor depletion and signaling inhibition, demonstrating the modularity of the platform. However, the optimal antibody geometry, binder stoichiometry, and TfR1 binder affinity varied across different targets. For instance, RXFP1 GTACs achieved the highest inhibition with a 2:1 RXFP1/TfR1 binder stoichiometry, whereas AT1R GTACs required bivalent TfR1 engagement. For CCR6, a 1:1 stoichiometry resulted in higher maximal depletion than a 2:2 stoichiometry. Additionally, low-affinity TfR1 binders were sufficient to achieve potent receptor depletion and signaling inhibition for AT1R and CCR6, while RXFP1 required high-affinity binding of both RXFP1 and TfR1 for high-potency antagonism. These findings underscore both the flexibility of GTAC designs and the importance of target-specific optimization to develop the most effective GTAC antagonists for each receptor.

## Discussion

In this study, we present, to our knowledge, the first comprehensive design and characterization of extracellular GPCR degraders, using five targets (AT1R, A2AR, APJ, RXFP1, and CCR6) and three detailed case studies (AT1R, RXFP1, and CCR6). GTACs consistently achieved high levels of target receptor depletion and degradation, with near-complete degradation observed in the case of CCR6. These results highlight GTACs as effective modalities for targeted degradation of GPCRs, with the potential to address previously undrugged receptors through targeted protein degradation. We evaluated these degraders not only in engineered HEK293 cell systems, which remain the predominant models for GPCR research, but also across native cellular contexts such as OVCAR-8 and Jurkat cells. These results demonstrated the broad applications of GTACs in both engineered and endogenous biological contexts.

Beyond demonstrating potent degradation activity, our work elucidates functional advantages of antibody-based degraders compared to conventional neutralizing antibodies. Most existing studies focus primarily on characterizing degradation efficacy, whereas the downstream consequences for receptor signaling remain less well characterized. Here, we present in-depth comparative analyses of the impacts of extracellular degrader constructs––versus antibody or Nb antagonists––on signaling outcomes. Our results show that degraders can achieve greater efficacy and improvements in antagonism potency by several orders of magnitude.

This is exemplified in our studies of RXFP1 and CCR6, two GPCR targets with significant therapeutic interest. In the case of RXFP1, the receptor’s extremely high ligand binding affinity and exquisite sensitivity to its ligand pose formidable obstacles to antagonism. Our Nb engineering efforts produced the first known RXFP1-targeting Nb antagonist. However, meaningful signaling inhibition was observed only at micromolar Nb concentrations in engineered HEK293 cells, and no significant effects were detected on downstream pro-tumorigenic gene expression in OVCAR-8 cells. In contrast, the RXFP1 GTAC achieved over 150-fold greater potency, delivering significant inhibition of downstream effectors. These results establish, for the first time, a biologics-based antagonist approach for RXFP1, demonstrating how degradation can enable functional inhibition where neutralization alone is insufficient.

Similarly, CCR6 is a chemokine receptor of considerable interest across inflammatory and autoimmune disease indications. However, the field has long struggled with insufficient inhibition of chemokine receptors due to allosteric activation mechanisms and high local chemokine concentrations. For CCR6, these mechanisms limit blocking antibodies to achieving partial reduction in signaling and cell migration. In contrast, our CCR6-targeting GTACs induced near-complete receptor degradation and fully blocked chemotaxis. These findings suggest a new strategy to modulate chemokine receptors, demonstrating that degraders can reach levels of pathway inhibition and phenotypic control not attainable with traditional antibody blockade.

Our work also advances the engineering principles of degrader modalities. Our multi-color imaging and small-molecule inhibitor analyses revealed that internalized receptors traffic to both recycling and late endosomes, and that protein synthesis inhibition reduces residual receptor levels. These results suggest a dynamic model in which internalization, degradation, recycling, and synthesis collectively determine receptor abundance under degrader treatment. We further discovered that while pH sensitivity in the GPCR-binding arm may not be essential for membrane protein depletion and signal antagonism, it may contribute to enhanced degradation. Additionally, we found that reducing TfR1 affinity can enhance effector recycling and, in certain cases, yield more potent and sustained receptor degradation and signaling inhibition. These mechanistic insights and protein engineering principles contribute novel understanding to degrader engineering and may have broad implications for designing and optimizing degrader constructs with similar architectures and target scopes.

In addition to therapeutic implications, our GTAC platform provides a valuable research tool for probing GPCR signaling and function. GTAC enables rapid on-demand, post-translational receptor depletion, allowing functional interrogation of GPCR pathways without relying on genetic knockout or knockdown approaches^65^. We demonstrated that a FLAG-targeting GTAC can achieve targeted cell surface depletion, receptor degradation, and signaling inhibition of GPCRs. Given the widespread use of FLAG-tagged GPCR constructs, including resources such as the PRESTO-Tango GPCR library^66^, this approach holds promise for rapid generalization and application to diverse receptors, even in cases where a target-specific binder is not available. In addition, GTACs provide a unique means to control GPCR localization from the cell surface to endolysosomal compartments, which may offer a new approach to dissect how the spatial origin of GPCR signals influences cellular outcomes. Notably, our findings show that GTACs can drive comparable levels of receptor internalization even for AT1R mutants with impaired β-arrestin binding. This offers an opportunity to investigate post-internalization signaling and trafficking behaviors of receptors carrying disease-relevant or functionally significant mutations. It is important to note, however, that in cases where the agonist is intrinsically membrane-permeable, as with the AZD5462 molecule in RXFP1 signaling, the GTAC strategy may be less effective as a purely inhibitory approach. In such contexts, GTACs instead may offer a powerful means to study the spatial dynamics of agonist activity at the cell surface versus within endolysosomal compartments.

Taken together, our findings position GTACs as a distinctive addition to the repertoire of strategies for GPCR modulation. While further studies will be required to define the translational scope of this approach across various GPCR classes and disease contexts, the principles demonstrated here establish a foundation for expanding extracellular targeted degradation as a tool for probing GPCR biology and addressing pharmacological challenges that have remained difficult to overcome with conventional modalities.

## Extended Data Figures

**Extended Data Figure 1.**
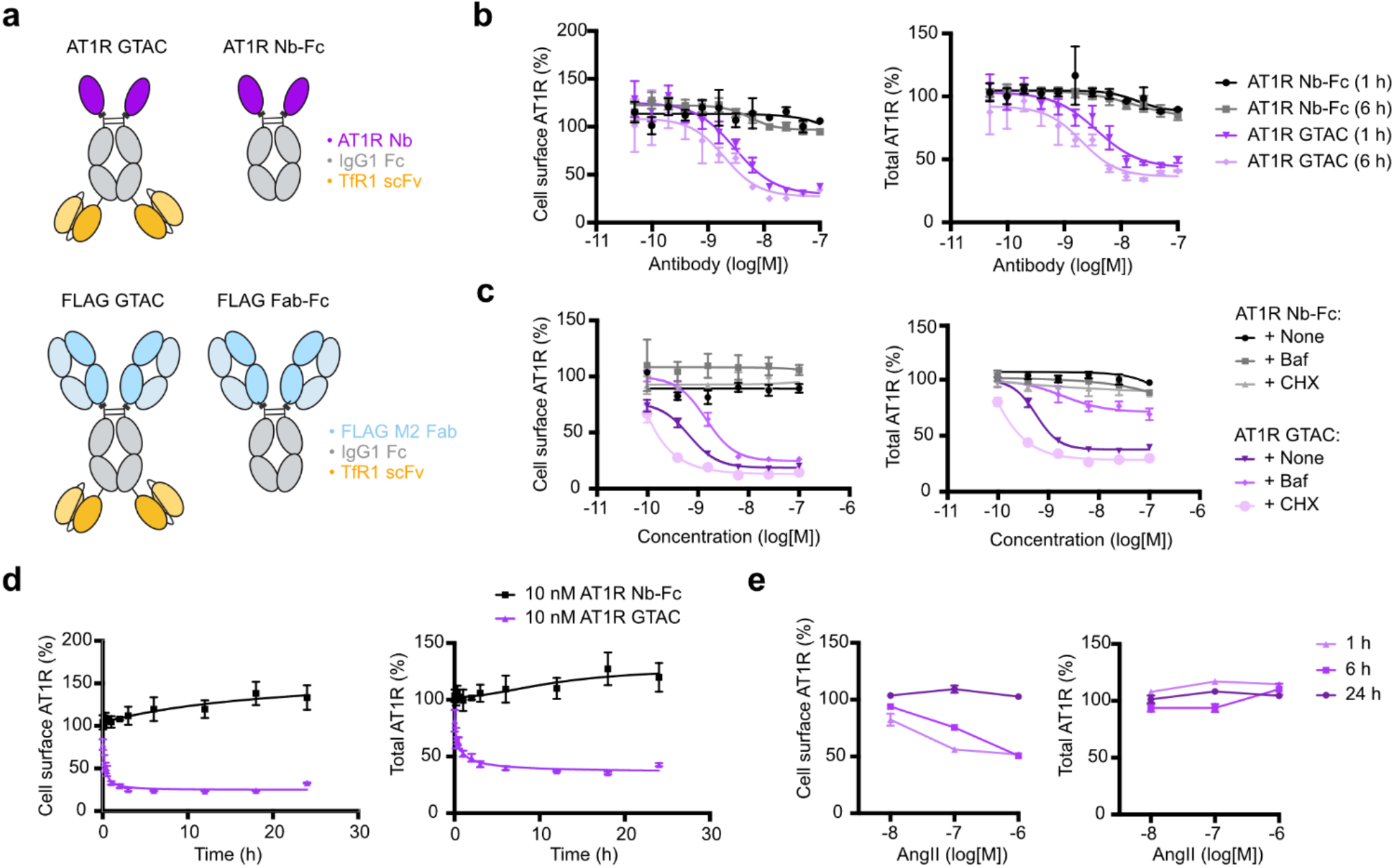
Kinetics of GTAC or AngII-mediated receptor internalization and degradation. **(a)** Schematic representations of AT1R GTAC, AT1R Nb-Fc, FLAG GTAC, and FLAG Fab-Fc constructs. **(b)** Cell surface AT1R levels (left, measured by anti-FLAG staining) and total AT1R levels (right, measured by GFP fluorescence) in FLAG-AT1R-GFP^+^ TfR1^+^ Expi293 cells after 1 h or 6 h treatment with varying concentrations of AT1R GTAC or Nb-Fc, assessed by flow cytometry. Data represent mean ± s.d. from n = 3 replicates. **(c)** Cell surface AT1R levels (left, measured by anti-FLAG) and total AT1R levels (right, measured by GFP) in FLAG-AT1R-GFP^+^ TfR1^+^ Expi293 cells after 6 h treatment with varying concentrations of AT1R GTAC or Nb-Fc, following 1.5 h pre-treatment with either 250 ng/mL bafilomycin A1 (Baf), 25 μg/mL cycloheximide (CHX), or no pre-treatment. Data represent mean ± s.d. from n = 3 replicates. **(d)** Cell surface AT1R levels (left, measured by anti-FLAG) and total AT1R levels (right, measured by GFP) in FLAG-AT1R-GFP^+^ TfR1^+^ Expi293 cells over a 24 h time course after treatment with 10 nM AT1R GTAC or 10 nM Nb-Fc. Data represent mean ± s.d. from n = 3 replicates. **(e)** Cell surface AT1R levels (left, measured by anti-FLAG) and total AT1R levels (right, measured by GFP) in FLAG-AT1R-GFP^+^ TfR1^+^ Expi293 cells following treatment with varying concentrations of AngII for 1 h, 6 h, or 24 h. Data represent mean ± s.d. from n = 3 replicates.

**Extended Data Figure 2.**
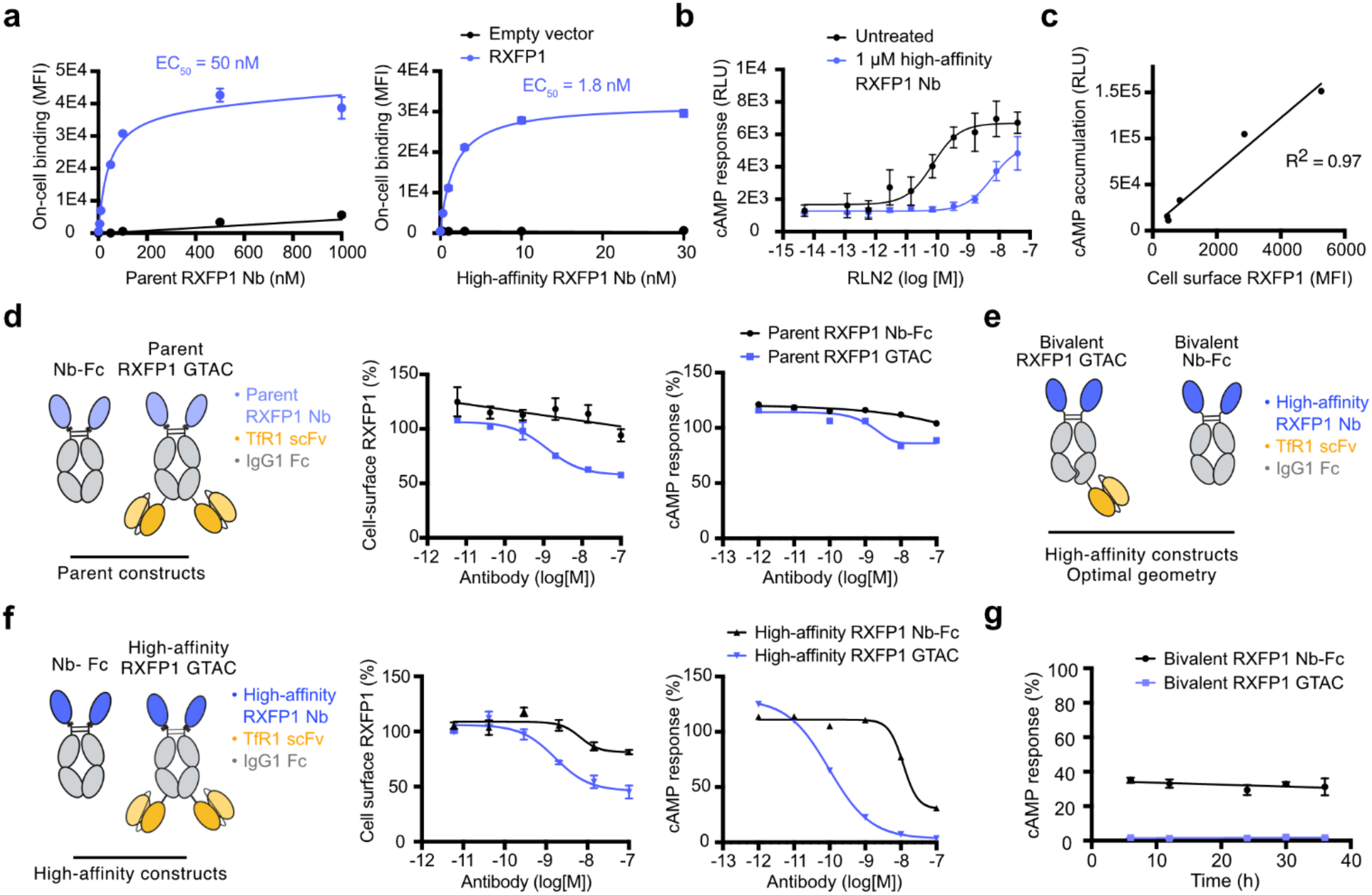
Potent RXFP1 antagonism with GTAC utilizing a high-affinity RXFP1 Nb. **(a)** On-cell binding of parent RXFP1 Nb and high-affinity RXFP1 Nb to Expi293 cells transfected with RXFP1 or empty vector, measured by flow cytometry. Data represent mean ± s.d. from n = 3 replicates. **(b)** Cellular cAMP levels in RXFP1^+^ GloSensor^+^ HEK293 cells measured by split-luciferase activity after 20 min pre-treatment with 0 or 1 μM high-affinity RXFP1 Nb, followed by 30 min stimulation with varying concentrations of RLN2. Data represent mean ± s.d. from n = 3 replicates. **(c)** Correlation between RXFP1 cell surface expression and RLN2-induced cAMP signaling. RXFP1 levels were titrated by 24 h induction with varying concentrations of dox, cell surface RXFP1 expression was measured by anti-FLAG flow cytometry, and cAMP response was measured by split-luciferase in FLAG-RXFP1^+^ TfR1^+^ GloSensor^+^ HEK293T cells after 30 min stimulation with 1 nM RLN2. Data represent mean ± standard deviation from n = 2 replicates. **(d)** Schematic representation of bivalent homodimeric RXFP1 GTAC and control Nb-Fc constructs utilizing the parent RXFP1 Nb (left). Cell surface RXFP1 levels (middle, measured by anti-FLAG) in FLAG-RXFP1^+^ TfR1^+^ Expi293 cells after 2 h treatment with GTAC or Nb-Fc. cAMP levels in RXFP1^+^ TfR1^+^ GloSensor^+^ HEK293T cells (right, measured by split-luciferase) after 2 h pre-treatment with GTAC or Nb-Fc, followed by 30 min stimulation with 1 nM RLN2. Data represent mean ± s.d. from n = 3 replicates (middle) and n = 1 replicate (right). **(e)** Schematic of optimized bivalent RXFP1 GTAC and Nb-Fc used in (g). **(f)** Schematic of bivalent homodimeric RXFP1 GTAC and control Nb-Fc constructs utilizing the high-affinity RXFP1 Nb (left). Cell surface RXFP1 levels (middle, measured by anti-FLAG) in FLAG-RXFP1^+^ TfR1^+^ Expi293 cells after 2 h treatment with GTAC or Nb-Fc. cAMP levels in RXFP1^+^ TfR1^+^ GloSensor^+^ HEK293T cells (right, measured by split-luciferase) after 2 h pre-treatment with GTAC or Nb-Fc, followed by 30 min stimulation with 1 nM RLN2. Data represent mean ± s.d. from n = 3 replicates (middle) and n = 1 replicate (right). **(g)** cAMP levels in RXFP1^+^ TfR1^+^ GloSensor^+^ HEK293T cells measured by split-luciferase, following pre-treatment with 10 nM optimized bivalent RXFP1 GTAC or Nb-Fc over 36 h and subsequent stimulation for 30 min with 1 nM RLN2. Data represent mean ± s.d. from n = 3 replicates.

**Extended Data Figure 3.**
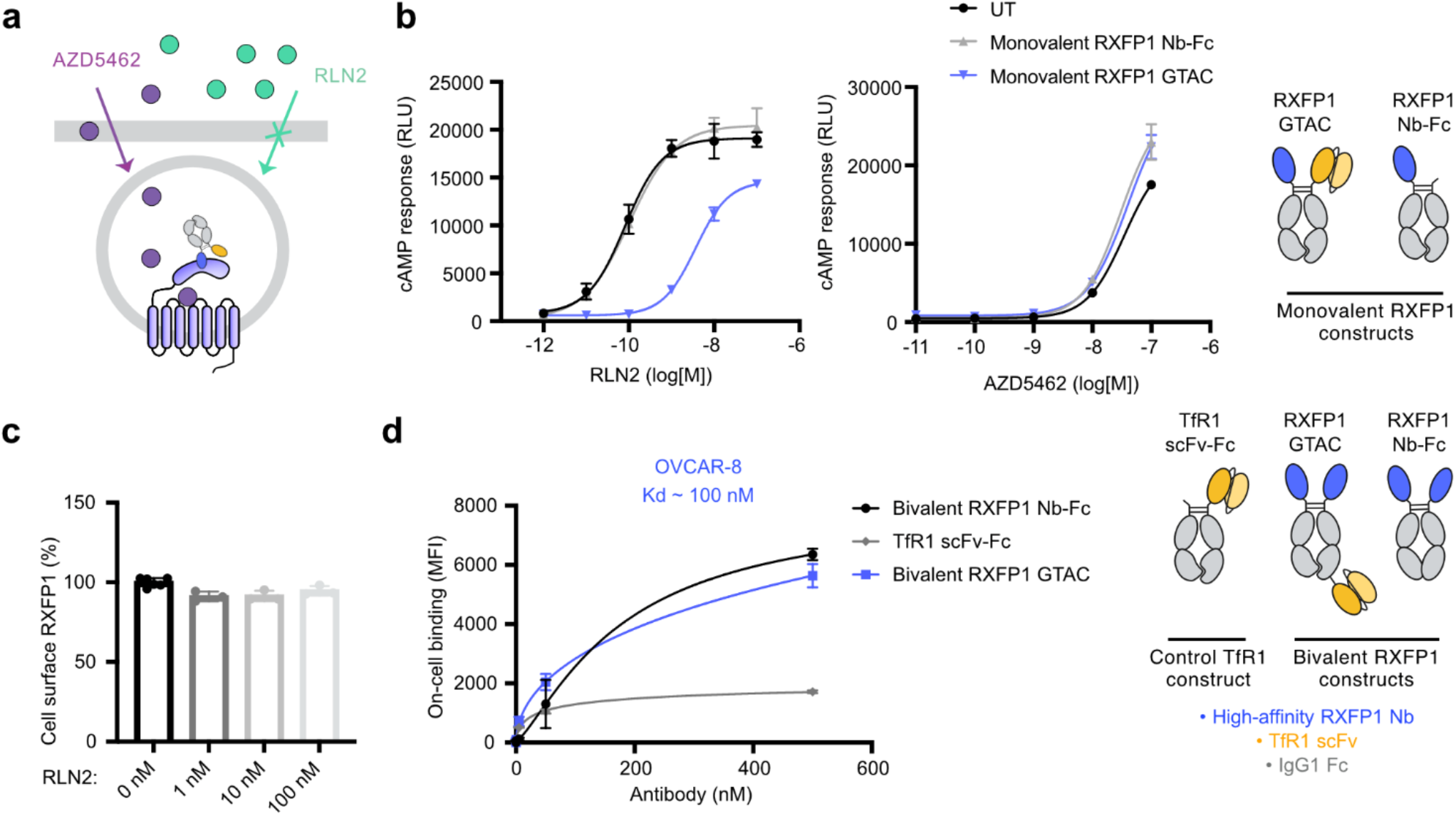
RXFP GTAC binding to OVCAR-8, and distinct GTAC responses with different ligands. **(a)** Schematic showing cell-permeant agonist AZD5462 versus impermeant native agonist RLN2. Unlike RLN2, AZD5462 may access RXFP1 after GTAC-induced internalization. **(b)** cAMP response measured by split-luciferase in RXFP1^+^ TfR1^+^ GloSensor^+^ HEK293T cells after 2 h pre-treatment with 10 nM monovalent RXFP1 GTAC or Nb-Fc, followed by 30 min stimulation with RLN2 (left) or AZD5462 (right). RXFP1 GTAC antagonized RLN2-but not AZD5462-induced cAMP response. Data represent mean ± standard deviation from n = 2 replicates. **(c)** Cell surface RXFP1 levels in FLAG-RXFP1^+^ TfR1^+^ Expi293 cells, measured by anti-FLAG flow cytometry after 6 h incubation with varying concentrations of RLN2. Data represent mean ± standard deviation from n = 3 replicates. **(d)** Bivalent RXFP1 Nb-Fc, GTAC, and TfR1 scFv-Fc binding to OVCAR-8 cells, measured by Protein A flow cytometry. Data represent mean ± standard deviation from n = 3 replicates.

**Extended Data Figure 4.**
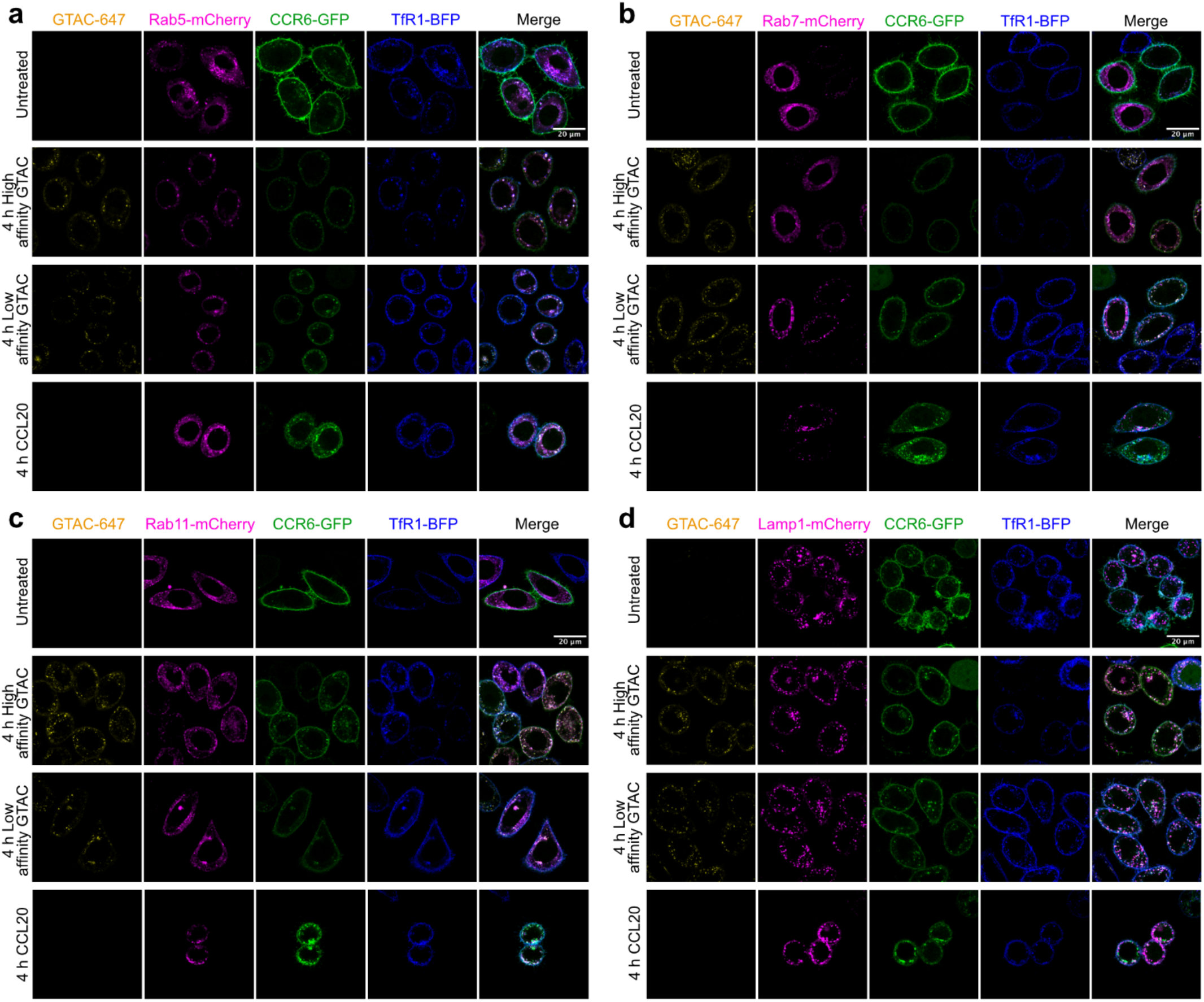
Intracellular trafficking of GTAC-mediated degradation of CCR6. Multiplexed four-color live-cell imaging in FLAG-CCR6-GFP^+^, TfR1-BFP^+^ HeLa cells expressing FLAG-CCR6-GFP and TfR1-BFP with 4 h treatment with 10 nM GTAC conjugated to Alexa Fluor 647 or CCL20. Colocalization is shown with mCherry-tagged endolysosomal markers including Rab5 **(a)**, Rab7 **(b)**, Rab11 **(c)**, or Lamp1 **(d)**, to indicate the early endosomes, late endosomes, recycling endosomes, or lysosomes, respectively. The high affinity GTAC is compared to the low affinity GTAC.

## Notes

### Competing Interest Statement

K.R., L.S., D.Z., and X.Z. have filed patent applications for the GTAC technologies. K.A. is a consultant to Odyssey Therapeutics, is on the SAB of CAMP4 Therapeutics, and received research funding from Novartis not related to this work. A.C.K. is a cofounder and consultant for Tectonic Therapeutic and Seismic Therapeutic, and for the Institute for Protein Innovation, a nonprofit research institute. X.Z. is a founder and consultant for VincenTx, a consultant for Merck, and receives research funding from Merck unrelated to this work.

### Funding

- National Institutes of Health: DP2GM154013, R00EB030587, R01CA260415, R01AR079489, R01AR081264, K99HD110612
- National Institutes of Health: R01GM134539, 2T32GM132089-06, T32GM139775
- Helen Gurley Brown Foundation
- DFCI Accelerator Fund
- Richard and Susan Smith Family Foundation
- Landry Cancer Biology Consortium
- Daiichi Sankyo Foundation of Life Science
- Japanese Dermatological Association
- Mochida Memorial Foundation for Medical and Pharmaceutical Research
- HHMI Hanna H. Gray Fellows Program

